# Contrast adaptation improves spatial integration

**DOI:** 10.1101/2021.03.14.435290

**Authors:** Noga Pinchuk-Yacobi, Dov Sagi

**Author notes:** Corresponding Author: Dov Sagi < >.

## Abstract

The effects of contrast adaptation and contrast area summation were investigated using a contrast discrimination task. The task consisted of a target of variable size, and a pedestal with a fixed base contrast. Discrimination performance was examined for a condition in which the pedestal size was fixed, equal to the largest target size, and for a condition in which the pedestal size matched the target size and thus varied with it. Repeated performance of the task produced rapid within-session improvements for both conditions. For stimuli with a matching size of target and pedestal, the performance improved only for the larger targets, indicating the development of area summation, which was initially absent for these stimuli. However, the improvements were mostly temporary, and were not fully retained between subsequent sessions. The temporary nature of the sensitivity gains implies that they resulted, at least in part, from rapid adaptation to the stimulus contrast. We suggest that adaptation decorrelates and thus reduces the spatial noise generated by a high-contrast pedestal, leading to improved area summation and better contrast sensitivity. A decorrelation model successfully predicted our experimental results.

## Introduction

Visual adaptation is a stimulus-driven process that continuously adjusts the neural responses to the statistics of the current visual environment. Contrary to perceptual learning, which leads to long-term improvements in performance, adaptation effects are generally short termed, tracking the current visual input to the eyes. However, both are thought to reflect neuronal plasticity, short-term in adaptation, and long-term in learning, possibly interacting (Sagi, 2011; Webster, 2015). One approach to visual pattern adaptation assumes that coding efficiency can be improved via short-term plasticity. According to this view, visual adaptation adjusts the limited dynamical range of individual neurons or a population of neurons to the statistics of the recent visual input (Wainwright, 1999; Wark et al., 2007). Coding efficiency can also be enhanced by reducing redundancy in sensory signals, possibly by decorrelating neuronal responses across processing channels (Barlow & Földiák, 1989). These continuous adjustments are expected to improve discriminability. However, previous studies that tested how adaptation affects contrast discrimination found only weak and mixed evidence for improved discriminability using target stimuli similar to the adapter (Barlow et al., 1976; Greenlee & Heitger, 1988; Määttänen & Koenderink, 1991a; Ross et al., 1993). In a recent study we found large improvements in a grouping discrimination task, which were attributed to visual adaptation (Pinchuk-Yacobi & Sagi, 2019). Perceptual grouping was suggested to rely highly on spatial correlations and spatial integration (Ben-Av & Sagi, 1995); therefore, it should be most affected by adaptation-induced spatial decorrelation. On the other hand, contrast discrimination, which is also influenced by spatial configuration (Adini & Sagi, 2001; Snowden & Hammett, 1998), does not benefit from spatial integration when the base contrast (pedestal) and the increment contrast (target) are over an equal spatial extent, explained by balanced excitation and inhibition interactions (Bonneh & Sagi, 1999b; Legge & Foley, 1980; Meese, 2004). However, spatial integration in contrast discrimination was observed for varying target sizes when a large, fixed, base contrast was used (Bonneh & Sagi, 1999b).

Here we aimed at testing the dynamics of contrast discrimination while varying target size, using stimuli in which the pedestal was either fixed at a maximal size, or of varied size, matching the size of the target. We tested for changes in observers’ performance, both repeatedly within a session as well as across several daily sessions. We found that repeated performance of the task produced rapid within-day improvements, which were largely transient, and were not fully retained when tested on subsequent days. For stimuli with a matching size of target and pedestal, the performance improved only for the larger targets, indicating the development of area summation, which was initially absent for these stimuli. We assumed that the transient nature of the gains implies that they resulted from rapid adaptation to the visual stimuli. Our results were predicted by a simple model, assuming that adaptation decorrelates spatial noise, thus leading to improved spatial integration and better contrast sensitivity. Practice with the task, over several days, resulted in faster adaptation, in accordance with previous findings (Yehezkel et al., 2010), supporting the idea that adaptation involves short-term plasticity, which with repetition, may consolidate and result in long-term effects.

## Methods

### Apparatus

The stimuli were presented on a 23.6’’ VIEWPixx/3D monitor (1920 × 1080, 10bit, 120Hz, with ‘scanning backlight mode’) viewed at a distance of 100 cm. The mean luminance of the display was 47.26 cd/m^2^, in an otherwise dark environment.

### Observers

Seventeen observers with normal or corrected-to-normal vision participated in the experiments described in the Results section. Eleven observers participated in the pilot experiment described in Appendix A, out of which one continued to the main experiments. All observers were naïve to the contrast discrimination task and gave their written informed consent. The work was carried out in accordance with the Code of Ethics of the World Medical Association (Declaration of Helsinki), and was approved by the Institutional Review Board (IRB) of the Weizmann Institute of Science.

### Stimuli and task

#### Contrast discrimination task (CD)

The stimuli consisted of two peripheral Gabor patterns (5.3° eccentricity left and right of fixation, 100 ms duration) presented simultaneously (Figure 1). Both patterns contained a pedestal signal (Gabor patch with λ = 0.53°; phase = 0) with a constant base contrast (Michelson contrast = 55%). One of the patterns, either on the left or on the right, contained an additional target signal (a Gabor patch with λ = 0.53°; phase = 0) that was added to the pedestal. The size of the target (defined as the standard deviation of the Gaussian envelope, σ) was one of four values: 0.37°, 0.56°, 0.84°, or 1.3°, which were randomly interleaved across trials. The size of the pedestal was either fixed at a maximal size (σ = 1.3°, ‘Non-matched’ condition) or changed in size in accordance with the size of the target (σ = 0.37°, 0.56°, 0.84°, or 1.3°, ‘Matched’ condition). Contrast thresholds were measured using a spatial two-alternative forced-choice (2AFC) paradigm. After fixating a central fixation circle, observers, when ready, initiated a new trial, and reported which of the patterns (left or right) contained the target, that is, which of the Gabor patterns appeared to have a higher contrast. An auditory feedback was given for an incorrect response.

**Fig. 1.**
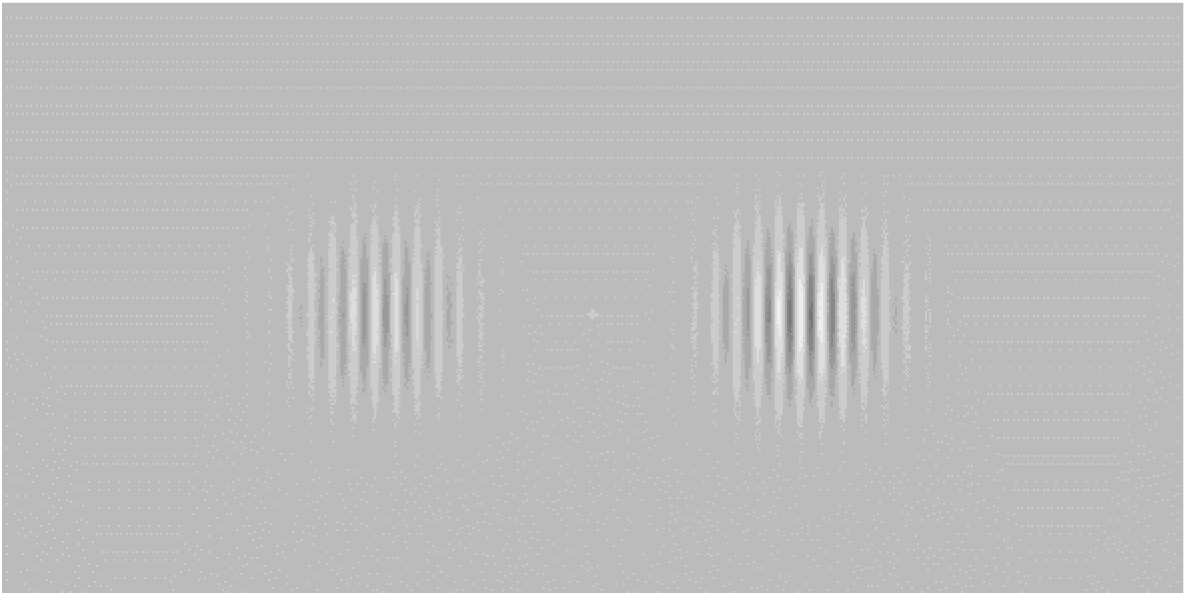
Stimuli and task. Observers fixated the white cross at the center of the screen and judged the location of a contrast increment (spatial 2AFC: left or right, 5.3° eccentricity, 100 ms duration).

### Procedure

Seventeen observers participated in this experiment. Ten of them participated in the ‘Non-matched’ condition, and seven participated in the ‘Matched’ condition. The experiment included one introductory session and four testing sessions, on different days. Each session consisted of four blocks with 140 or 168 trials per block (4 target sizes, × 7 contrast values, and × 5 or 6 trials per contrast). The target contrast was one of seven values: 0.8%, 1.6%, 3.1%, 4.7%, 6.3%, 12.6%, and 25.2%, randomly intermixed across trials. The measured psychometric curves (the percentage of correct responses as a function of target contrast) were fitted with a Weibull function distribution (lapse rate: up to 0.05). The discrimination threshold was defined as the contrast for which there were 79% correct responses. Fitting was performed using Psignifit 4.0 software for MATLAB (Schütt et al., 2016).

## Results

Results for the group contrast discrimination thresholds (averaged across observers) for the ‘Non-matched’ group (fixed pedestal size) and the ‘Matched’ (varied pedestal size) groups are presented in Figure 2A and Figure 2B, respectively. To test for significant within-session improvements, we ran a repeated measures ANOVA with the within-observer factors including day (4 days), size (4 sizes), and block (first block vs last block) performed separately for the two groups. Performance within-session improved significantly in both groups, as indicated by a significant decrease in the threshold from the first block to the last block (5.2±0.8, F(1,9) = 42.8, p< 0.001; 2.2±0.6, F(1,6) = 11.6, p =0.01, mean±SEM in the ‘Non-matched’ group and the ‘Matched’ group, respectively). In the ‘Non-matched’ group, there was also a significant interaction between block threshold and day (F(3,9) = 3.4, p< 0.05). To determine how the within-session improvements changed between days, we ran separate ANOVAs for each daily session. Results for the ‘Non-matched’ group show significant within-session improvements in every daily session (8.7±2.1, F(1,9)=17.1, p<0.01, 4.0±0.8, F(1,9)=22.4, p=0.001, 3.5±1.2, F(1,9)=8.2, p<0.05, 4.4±0.9, F(1,9)=23.8, p=0.001, improvements within the 1^st^, 2^nd^, 3^rd^, and 4^th^ daily sessions, respectively). However, in the ‘Matched’ group, only day 3 reached significant within-session improvement (2.1±1.5, F(1,6)=2.0, p=0.2; 1.8±1.0, F(1,6)=3.5, p=0.1; 3.2±1.3, F(1,6)=6.3, p<0.05; 1.6±0.7, F(1,6)=5.7, p=0.06, mean±SEM, improvements within the 1^st^, 2^nd^, 3^rd^, and 4^th^ daily sessions, respectively). To determine how the within-session improvements changed between sizes, we ran separate ANOVAs for each stimulus size. Results for the ‘Non-matched’ group show significant within-session improvements for all sizes (6.0±1.3, F(1,9)=21.0, p=0.001, 7.1±1.9, F(1,9)=13.9, p<0.01, 4.3±0.9, F(1,9)=23.5, p=0.001, 3.2±1.4, F(1,9)=5.2, p=0.05, improvements in the 1^st^, 2^nd^, 3^rd^, and 4^th^ target sizes, respectively). However, in the ‘Matched’ group, only the two large target sizes (sizes 0.84° and 1.3°) reached significant within-session improvement (2.8±1.7, F(1,5)=2.9, p=0.1; 0.8±0.9, F(1,5)=0.7, p=0.4; 2.0±0.7, F(1,5)=8.4, p<0.05; 3.1±1.0, F(1,5)=9.4, p<0.05, improvements in the 1^st^, 2^nd^, 3^rd^, and 4^th^ target sizes, respectively).

**Fig. 2.**
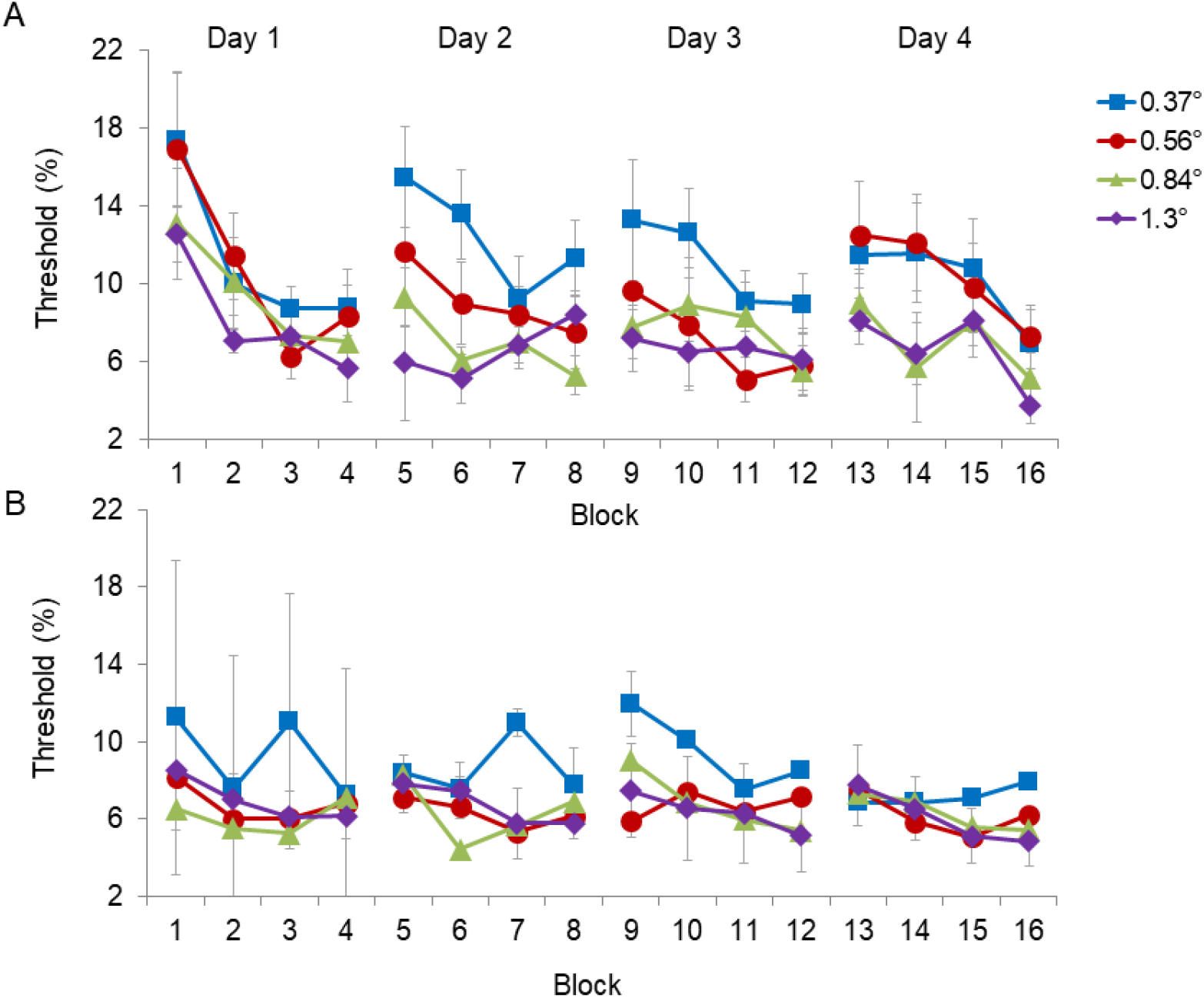
Contrast discrimination thresholds for stimulus configurations in which a target Gabor pattern (with sizes: 0.37°, 0.56°, 0.84°, or 1.3°) was added to a pedestal Gabor pattern that was either with a fixed maximal size (A, ‘Non-matched’ group), or with a size equal to the target size (B,‘Matched’ group). Results show within-day improvements that are not fully retained in subsequent daily sessions. Thresholds represent averages across observers (A: N = 10, B: N=7), with error bars corresponding to ±1 standard error of the mean.

Figure 2 shows that most of the within-session gains did not survive the time between sessions: the first measured threshold during a session was higher than the last measured threshold in a previous session. To further test the perseverance of the within-session improvements, we ran a repeated measures ANOVA with the within-observer factors of day (day 2-day 4), size (4 sizes), and block (the last block of the previous session vs the first block of the next session). In both groups the performance significantly deteriorated from the end of the previous session to the beginning of the next session (3.4±0.8, F(1,9)=17.8, p<0.01; 1.9±0.8, F(1,9)=5.9, p=0.05 in the ‘Non-matched’ group and the ‘Matched’ group, respectively), showing that the within-session gains were not fully retained between sessions. No significant interactions were found in this ANOVA analysis. This effect suggests that adaptation processes rather than learning played a role here (Sagi, 2011).

Next, we tested for between-session improvements by running a repeated measures ANOVA with only the thresholds of the first day (day1) and the last day (day 4). In both groups, there was no significant change in the overall daily performance (including all blocks of trials in a daily session) from the first session to the last session (1.2±1.1, F(1,9) = 1.3, p=0.3, 0.7±0.5, F(1,6) = 1.7, p=0.3 in the ‘Non-matched’ group and the ‘Matched’ group, respectively). However, in the ‘Non-matched’ group, there was a significant interaction between day and block (Day*Block), indicating that learning was different for different blocks (F(1,9) = 6.0, p<0.01). Testing for learning effects separately at the start (block 1) and at the end (block 4) of the session showed that significant learning occurred only at the start of the session, and only for the ‘Non-matched’ group (start: 6.1±1.6, F(1,9) = 14.6, p <0.01, 1.3±0.8, F(1,6) = 2.4, p =0.2; end: 1.9±1.5, F(1,9) = 1.6, p =0.2, 0.8±1.2, F(1,6) = 0.4, p =0.6 in the ‘Non-matched’ group and the ‘Matched’ group, respectively).

To directly compare the groups (Figure 3), we added to the repeated measures ANOVAs the between-observer factor of group type (‘Non-matched’ group or ‘Matched’ group). At the beginning of the session (first block), the performance of the ‘Matched’ group was significantly better than the performance of the ‘Non-Matched’ group (4.1±1.8, F(1,15) = 5.1, p <0.05). However, during the session, the performance differences between the groups diminished, and they were no longer significant at the end of the session (1.2±1.0, F(1,15) = 1.2, p=0.3). The significantly better performance of the ‘Matched’ group at the beginning of the session was found to depend on the day (significant interaction, F(3,15) = 4.0, p=0.01). Separate analyses for each day showed significant effects on all days except on day 3 (8.0±2.8, F(1,15)=8.2, p<0.01, 4.5±2.1, F(1,15)= 4.7, p<0.05, 0.8±2.4, F(1,15)=0.1, p=0.7, 3.2±1.3, F(1,15)=5.8, p<0.05, performance differences at the beginning of daily sessions, the 1^st^, 2^nd^, 3^rd^, and 4^th^, respectively). Apparently these differences between the groups, at the beginning of the session, decreased from day to day, probably due to the significant between day (first day vs last day) improvement at the beginning of the session (first block) only in the ‘Non-Matched’ group (previously reported), which was significantly higher than the corresponding insignificant improvement in the ‘Matched’ group (F(1,15) = 5.5, p< 0.05). In addition, the within-session improvements were significantly higher in the ‘Non-Matched’ group (F(1,15) = 7.6, p= 0.01), probably due to the significantly higher thresholds in the ‘Non-matched’ group at the beginning of the sessions, since there were no significant threshold differences at the end of the sessions. The deterioration between sessions (F(1,15) = 1.5, p= 0.2), and the threshold differences between target sizes at the beginning of the session (F(3,15) = 2.1, p= 0.1) and at the end of the session (F(3,15) = 0.5, p= 0.7) were not significantly different between the groups.

**Fig. 3.**
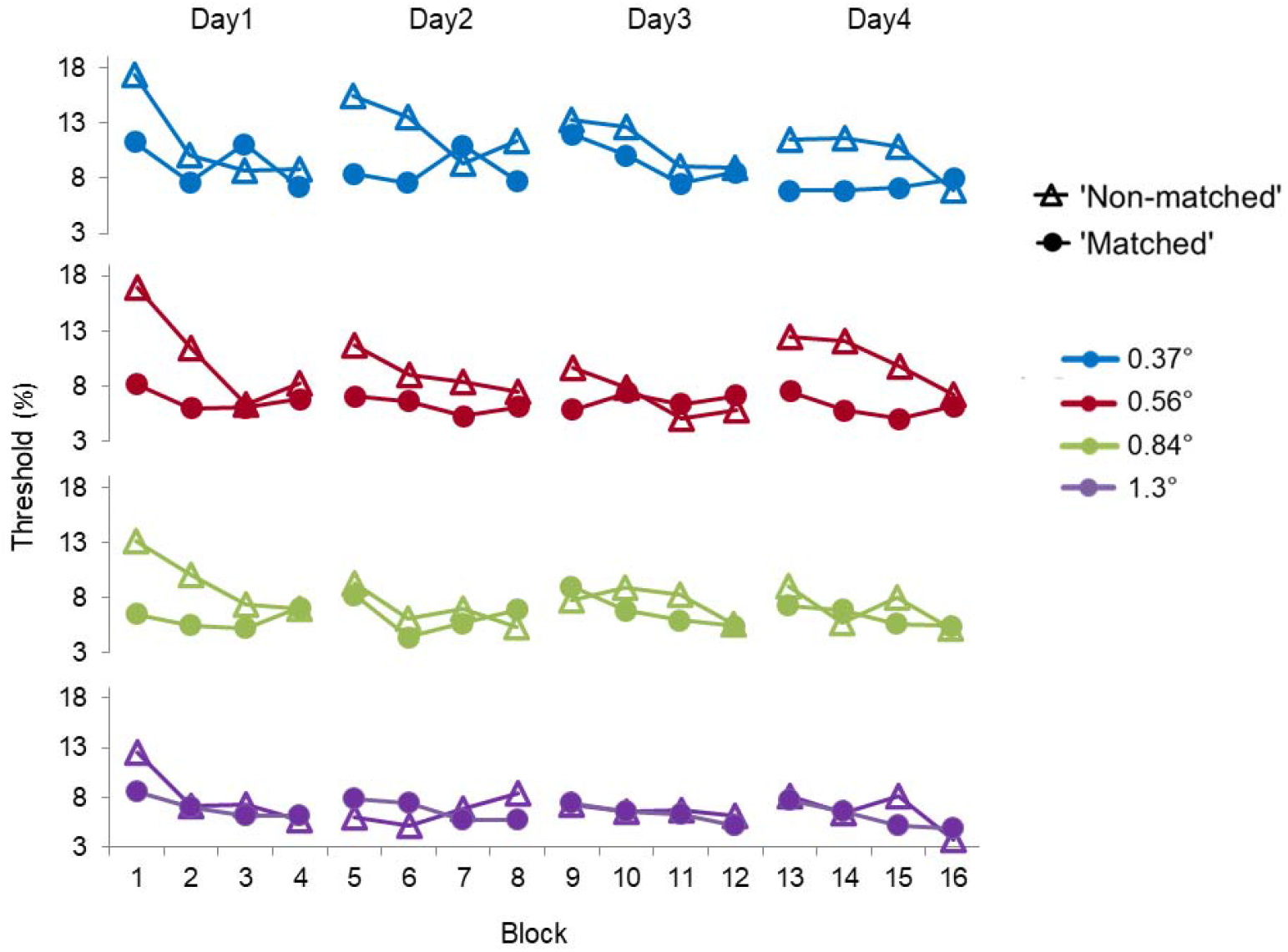
Comparing discrimination thresholds in the ‘Non-matched’ group (triangular markers) vs the ‘Matched’ group (circular markets) for different target sizes. At the beginning of each daily session the thresholds in the ‘Non-matched’ group were generally higher than those in the ‘Matched’ group, especially for the smaller target sizes (red and blue markers). However, thresholds at the end of the sessions (560 or 672 trials) were equivalent in both groups. Thresholds represent averages across observers (N = 10, N=7, for the ‘Non-matched’ and ‘Matched’ groups, respectively). Note that for the largest size (1.3°) the ‘Matched’ and the ‘Non-matched’ stimuli are identical but are presented within different mixtures of trials.

One of the main objectives of the current study was to test the effect of contrast adaptation on contrast area summation. Improved performance with increasing target size suggests that area summation exists in the contrast discrimination process. Figure 4 shows that at the beginning of the session the performance was significantly enhanced with target size only in the ‘Non-matched’ group (F(3,9) = 4.5, p =0.01, F(3,6) = 2.3, p =0.1, the ‘Non-matched’ group, and the ‘Matched’ group, respectively), indicating significant area summation only in the ‘Non-matched’ group. However, at the end of the session the performance in both groups was significantly enhanced with increasing target size (F(3,9) = 4.1, p <0.05, F(3,6) = 5.0, p =0.01 for the ‘Non-matched’ group and the ‘Matched’ group, respectively), indicating that area summation also developed in the ‘Matched’ group. These differences between groups did not reach significance (F(3,15)=2.1, p=0.1, F(3,15)=0.5, p=0.7, at the start of a session (1^st^ block) and at the end of a session (4^th^ block), respectively). To quantify the extent of the area summation, we calculated the summation slope of the (log) average threshold (across days and observers) as a function of the (log) squared value of the target size (see Figure 4). We then tested how the summation slope changes from the start of the session (block 1) to the end of the session (block 4). For the ‘Non-matched’ group the summation slopes were significantly negative at the start and at the end of the session (−0.23±0.07, p=0.01, paired t-test) (−0.18±0.06, p=0.01, paired t-test), indicating significant area summation throughout the session. For the ‘Matched’ group the summation slopes at the start of the session were not significantly different from zero (−0.05±0.07, p=0.5, paired t-test), indicating the lack of area summation. However, the summation factor at the end of the session became significantly negative (−0.14±0.03, p= 0.001, paired t-test), indicating the development of area summation in the ‘Matched’ group.

**Figure 4:**
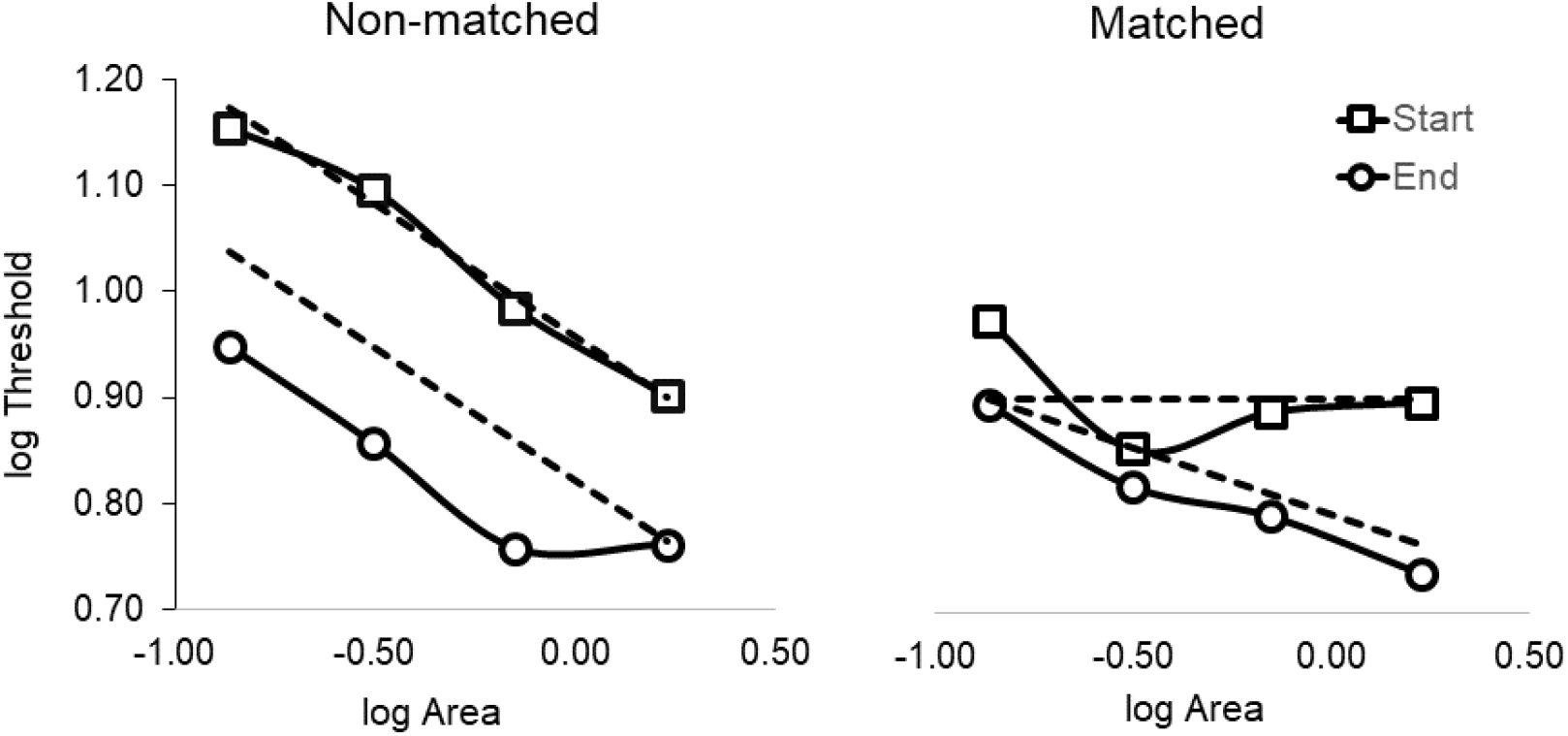
Contrast area summation at the start and at the end of a session for the Non-Matched and Matched conditions. ‘Start’ and ‘End’ results are from the first and the last blocks of trials in each session, respectively, averaged across all days (N=4) and observers (N=10 & 7 for the Non-Matched and the Matched conditions, respectively). The ‘Start’ results confirm previous results, showing that thresholds are roughly independent of stimulus area in the Matched condition; however, they improved with increasing area in the Non-Matched conditions, following the 4^th^ root summation rule (Bonneh & Sagi, 1998). The ‘End’ results show that area summation was developed in the ‘Matched’ condition during the session. Dashed lines denote the predictions of a model, assuming the 4^th^ root summation rule with spatially correlated noise at the ‘Start’ of the session and uncorrelated noise at the ‘End’ of the session (see the text for details).

Table 1 summarizes the statistic results.

**Table 1.**
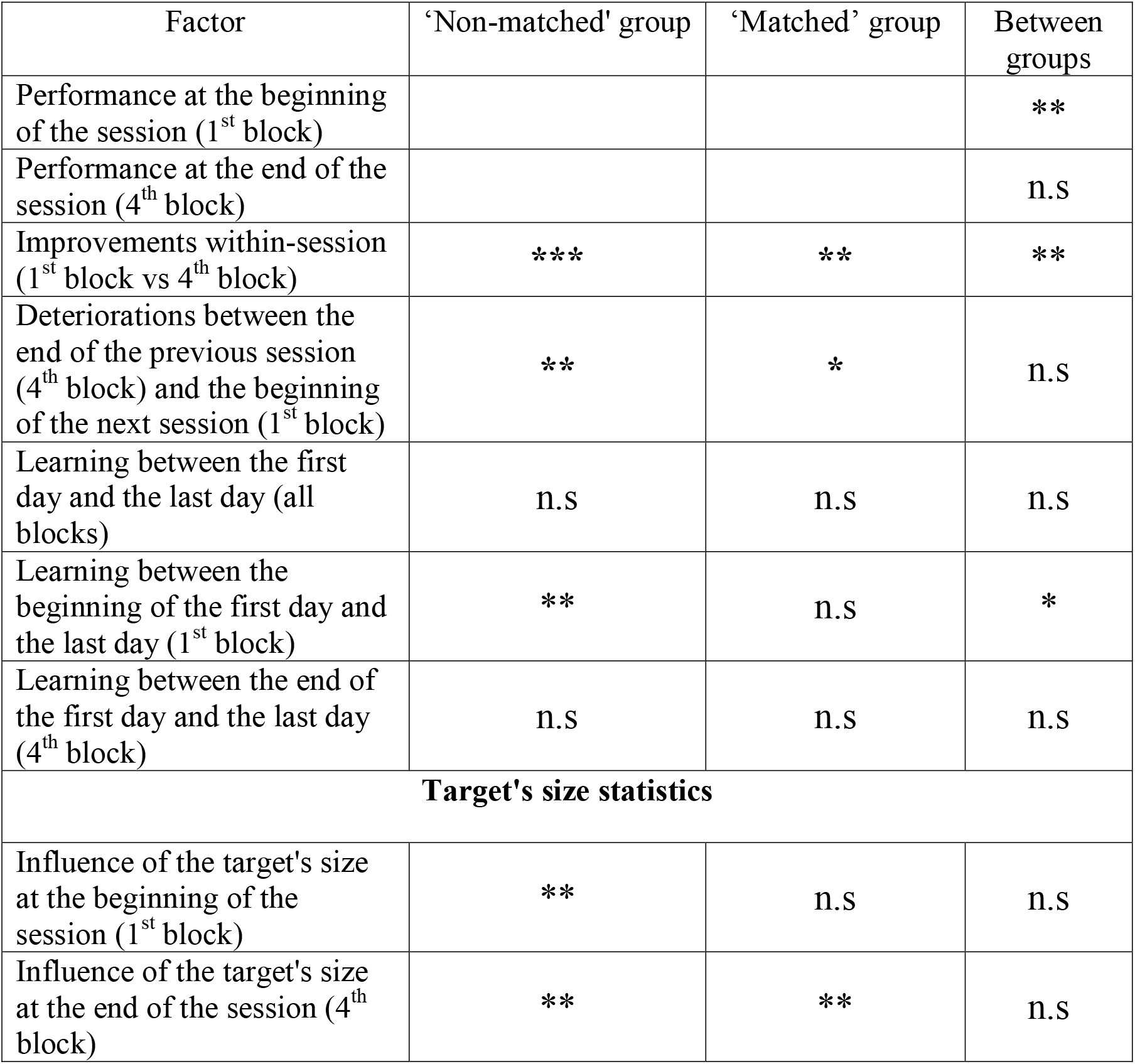
Statistical results for Exp2, ‘Matched’ vs ‘Non-matched’ conditions. *P < 0.05, **P < 0.01,***P < 0.001.

### The decorrelation model

To explain our results for the contrast discrimination thresholds and explain how thresholds change with the adaptation state, we constructed a simple toy model, based on the following assumptions:

1. Each stimulus (Left and Right) is processed by multiple functional units covering the stimulus area, creating two activity maps. Receptive fields of individual units are assumed to match the smallest target size (Gabor shaped, σ=0.37°, λ=0.53°).
2. Spatial integration of signal and noise:
Following Quick (1974), we defined the task relevant signal (*S*) to be the integrated incremental contrast

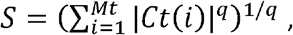

where *M_t_* is the number of functional stimulus units covering the target, *C_t_(i)* is the local incremental contrast contributed by the target, and *q=4*, assuming the well-documented 4^th^ root summation rule (Bonneh & Sagi, 1998; Graham, 1989; Meese & Summers, 2012; Usher et al., 1999). Here, following Quick (1974), we interpreted this equation as a nonlinear response to pooling prior to noise application, rather than a probabilistic summation (Graham, 1989), based on results showing that spatial integration depends on orientation and spatial proximity (Bonneh & Sagi, 1998).From the above equation, it follows that target area summation (*A_t_*) can be expressed as

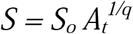

We assume the presence of target-independent response noise integrated across the pedestal area (e.g., on the non-target side). We formalized this integration using the above Quick (1974) equation, applied to the equivalent external noise. Thus, for correlated noise the integrated noise amplitude (*N*) across the pedestal area (*A_p_*) is

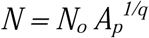

And for uncorrelated noise it is

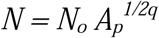
3. Noise correlations:
The noise is assumed to be spatially correlated before adaptation (block 1), but spatially uncorrelated after adaptation (block 4).

Based on the above assumptions, we estimated the dependence of target sensitivity (*S/N*) on the stimulus area before and after adaptation. The target strength (*S*) is proportional to the target_area^1/*q*^ independent of the adaptation state and the experimental condition. The noise level (*N*) depends on the pedestal size and on the state of adaptation:

In the Non-Matched condition, we assume that the noise is independent of the target area, set by the fixed pedestal extent (note that the different target sizes are presented in random order within each block of trials; thus, the integration area cannot be adjusted to the target size, but see below). Thus,

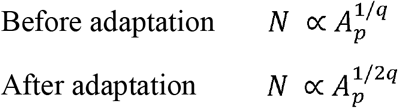

Therefore, 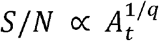, before and after adaptation.

In the Matched condition, the noise depends on the target area, since the pedestal area matches the target area:

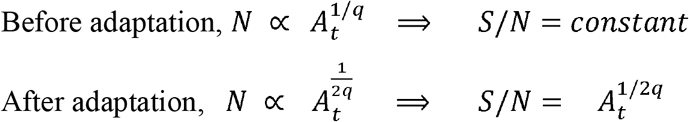

### Predictions

Slopes of log(threshold) vs log(target_area) are −0.25 in the ‘Non-matched’ condition, before and after adaptation, and 0 and −0.125 in the ‘Matched’ condition, before and after adaptation, respectively (Figure 4, dashed lines).

### Model fitting results

To test our model predictions, we plotted the measured thresholds as a function of the target area (σ^2^) using a log-log scale for both the start and the end of the sessions (Figure 4). To match the model predictions with the data, we used the ‘Start’ threshold of the largest target as a reference, averaging the corresponding measurements from the two conditions (these particular thresholds are available from both experimental groups and are expected to be equal at the beginning of the session, as indeed measured). Other thresholds are derived using the fourth root summation rule, assuming adaptation-dependent noise correlation (fully correlated before adaptation and limited to the receptive field size after adaptation). As shown in Figure 4, the predicted thresholds obtained from the model are very similar to the measured thresholds. At the beginning of the session, there was a significant area summation in the ‘Non-matched’ condition, with a (log-log) summation slope of −0.24, close to the predicted slope of −0.25. In the ‘Matched’ condition the summation slope was −0.05, compared with 0 predicted. However, at the end of the session, there was significant area summation in both conditions. In the ‘Non-matched’ condition, the value of the summation slope was −0.18 (but see below), compared with −0.25 predicted, and in the ‘Matched’ condition the value was −0.14, compared with −0.125 predicted.

The model overestimates the adapted thresholds. This can result from our assumption that *S* and *N* are integrated over all of the pedestal area. This assumption is motivated by the uncertainty imposed on the observer regarding the target size. However, it is possible that, during the experimental session, observers narrow down the integration area to reduce noise, thus improving sensitivity with smaller targets, while losing it with larger ones. This hypothesis is supported by the results showing no area summation with the largest target size after adaptation in the ‘Non-Matched’ condition (note that without this datum point the area summation slope is −0.27). Additionally, the predicted post-adaptation thresholds depend on the assumed receptive field size, here set to the smallest target size. Smaller receptive fields will increase the decorrelation gain in both experimental conditions.

## Discussion

In the experiment described here, observers performed a contrast discrimination task with a fixed base contrast (pedestal), and a variable target size. We compared discrimination behavior between an experimental condition, in which the pedestal size was fixed, equal to the largest target size (‘Non-matched’ condition), and a condition in which the pedestal size matched the target size and thus varied with it (‘Matched’ condition). Discrimination performance significantly improved during the sessions for both conditions. In the ‘Non-matched’ condition, the performance improved for all target sizes. In the ‘Matched’ condition performance improved only for the larger targets, indicating improvement by means of area summation. However, these within-session improvements were mostly temporary and were not fully retained between sessions (Figs. 2 and 3). The transient nature of these improvements implies that they result, at least in part, from an adaptation process that rapidly adjusts the visual system to the current statistics of the visual stimulation. Theoretical considerations suggest that these adjustments reduce input redundancy by a decorrelation process. Such a process is thought to improve coding efficiency (Barlow & Földiák, 1989). Accordingly, we suggest that adaptation improves discrimination performance as well as area summation by decorrelating and thus reducing the background noise generated by the high-contrast stimulus (i.e., the pedestal). A decorrelation analysis of our experiments was found to successfully predict the results (Figure 4), as described in the results section.

At the beginning of the sessions, in agreement with previous studies (Bonneh & Sagi, 1999b), there were significant differences between the conditions, regarding the general performance level and regarding how performance changed with target size (see Figure 3 and 4 thresholds at the start of the sessions). In the ‘Non-matched’ condition, with fixed pedestal size, increasing the target’s size reduced thresholds, indicating area summation. The (log(Threshold) vs log(Area)) summation slope was around −1/4 (fourth-root summation), in agreement with contrast discrimination studies (Bonneh & Sagi, 1999b), and with studies of contrast detection (Robson & Graham, 1981). However, in the ‘Matched’ condition, in which the pedestal size matched the target size, performance at the beginning of the session was almost equal for all the sizes (summation slope = −0.05).

These results are in agreement with previous studies showing that area summation is much reduced, practically non-existent, for a suprathreshold pedestal whose size matches the target’s size (Bonneh & Sagi, 1999b; Legge & Foley, 1980; Meese, 2004). Performance was significantly better in the =Matched’ condition, compared with the ‘Non-matched’ condition for the smaller targets. This was due to the larger pedestal in the ‘Non-matched’ condition, which extended beyond the target area (except for the largest target size), thus forming an extra surround that reduced the target’s visibility. Such a surround suppression was previously shown (Bonneh & Sagi, 1999b; Meese, 2004; Meese et al., 2005), and is consistent with studies showing reduced contrast discrimination performance with a high-contrast pedestal in the presence of an iso-oriented surround (Adini & Sagi, 2001; Chen & Tyler, 2001; Chen & Tyler, 2008, Bonneh & Sagi, 1999a; Snowden & Hammett, 1998; Wilkinson et al., 1997; Xing & Heeger, 2000; Zenger-Landolt & Koch, 2001), especially at the periphery (Petrov et al., 2005; Wilkinson et al., 1997; Xing & Heeger, 2000), supporting the involvement of spatial lateral-inhibitory interactions (Polat & Sagi, 1993; Sagi & Hochstein, 1985).

In contrast with the beginning, at the end of the sessions the differences between the conditions vanished. A moderate area summation developed in the ‘Matched’ condition, in which discrimination thresholds improved only for the larger targets, resulting in a significant area summation in both conditions (see Figure 4, thresholds at the end of the sessions). In the ‘Non-matched’ condition, performance significantly improved during the session for all target sizes, resulting in equivalent performance levels in both conditions (see Figure 3, thresholds at the end of the sessions). These discrimination improvements were mostly not retained between sessions (but see below); thus, they are considered here to result from contrast adaptation. Within the context of the standard contrast discrimination models and assuming contrast gain control (Foley, 1994; Foley & Chen, 1999; Meese, 2004) by a normalization process (Carandini & Heeger, 2012), our results can be implemented as dynamic normalization (Ross & Speed, 1991). According to these models, the response to each stimulus (either target+pedestal or pedestal alone) is proportional to the integrated excitatory inputs from all local units activated, divided by the activity in a ‘normalization pool’. The ‘normalization pool’ consists of a weighted sum of all units’ responses, activated by the target stimulus, the pedestal, and other contextual elements. In this type of models, adaptation can improve discrimination performance in the ‘Non-matched’ condition for all target sizes, by reducing the suppressive effects produced by the large and fixed pedestal. Weakening of surround suppression signals by adaptation is supported by electro-physiology (Patterson et al., 2013, 2014; Wissig & Kohn, 2012), especially after a prolonged adaptation time (Patterson et al., 2013). The development of area summation in the ‘Matched’ condition can result from asymmetric changes in the excitatory and suppressive summation weights due to adaptation. This suggestion is supported by results showing that adaptation in the barrel cortex shifts the balance between excitation to inhibition toward excitation, by adapting the inhibitory inputs more than the excitatory inputs (Heiss et al., 2008). In agreement with our results, recording from macaque V1 shows that by suppressing the neurons’ surround more than the neurons’ center, adaptation changes the balance of center and surround gains. Such a change of balance was shown to increases the neurons’ receptive field and consequently, increased the cortical summation (Cavanaugh et al., 2002). Functionally, surround suppression and normalization processes are thought to remove local correlations in neuronal activity arising from redundancy within natural images (Schwartz & Simoncelli, 2001). Barlow (1990) suggested adaptation as a process by which such inhibitory processes are strengthened to reduce correlations, an idea supported by our modeling results. Strengthening of inhibitory connections during adaptation may reduce the saliency of uniform stimuli, thus increase detection thresholds (Ross & Speed, 1991), and improve detection of texture boundaries (Sagi, 1995).

Previous psychophysical studies that tested how adaptation affects contrast discrimination found only weak and inconsistent evidence of improved discriminability for stimuli similar to the adapter (Abbonizio et al., 2003; Barlow et al., 1976; Foley & Chen, 1997; Greenlee & Heitger, 1988; Määttänen & Koenderink, 1991a; Wilson & Humanski, 1993). The adaptation effects observed here were larger in the ‘Non-Matched’ condition using stimuli that are quite different from those used in traditional contrast discrimination studies. Previous studies used different methods, but all had the target and the pedestal of an equal area, as in our ‘Matched’ condition. We note that the ‘Matched’ effects are relatively small and are significant only with the larger stimuli. Indeed, Greenlee & Heitger (1988), who used high-contrast (0.8) gratings, found that adaptation improved the discriminability of high-contrast targets. In contrast, Foley & Boynton (1993), Määttänen & Koenderink (1991b), and Wilson & Humanski (1993), who used narrowly localized Gabor stimuli, found that adaptation either did not affect the discrimination of high base contrast (Foley & Boynton, 1993), or only slightly improved discrimination in the highest base contrasts, but not for all observers (Wilson & Humanski, 1993). Interestingly, for the observer who initially did not improve in the Wilson & Humanski (1993) experiment, increasing the stimulus spatial frequency, which allowed for more area summation, resulted in some discrimination enhancement with a high base contrast. Barlow et al. (1976), using grating stimuli, found that adaptation had no effect on contrast discrimination at high base contrasts. However, Barlow measured discrimination either by using vertically split-field contrasts, which required only local discrimination around the vertical central line, or by detecting the point in time when a change in contrast occurred, which can be viewed as a change-detection task. Importantly, stimuli presented in the fovea benefit less from spatial integration due to the large sensitivity drop around the fovea. In our experiments, stimuli were presented in the near periphery, activating areas of relatively uniform sensitivity. Notably, unlike other studies using passive adaptation, in our experiments the adapted stimuli were task related, and the observers were asked to make an explicit judgment regarding each of them. Previous studies showed that passive adaptation is sufficient to cause adaptation-dependent sensitivity improvements (Greenlee & Heitger, 1988; Pinchuk-Yacobi & Sagi, 2019). However, top-down influences such as task relevance and attention can enhance the effects for some types of adaptation but not for others (Festman & Ahissar, 2004; Pinchuk-Yacobi et al., 2016).

Comparing the within-session dynamics in our study to other contrast discrimination studies is difficult, since nearly none reported performance changes within sessions, only the session means. The only study we found that did report within-session dynamics is a study by Yu et al. (2004), which supported our results of improved performance within sessions, which was not preserved between sessions.

Recent studies (Censor & Sagi, 2008; Pinchuk-Yacobi et al., 2016; Yehezkel et al., 2010) have suggested that an interaction exists between sensory adaptation and perceptual learning. Whereas adaptation is probed by temporal within-session changes, learning is thought to improve visual sensitivity across sessions. Here we show a significant across-session improvement in performance at the start of the session in the ‘Non-matched’ condition, but not in the ‘Matched’ condition. This across-session learning in the ‘Non-matched’ group can be compared to the learning found in contrast discrimination with flankers (Adini et al., 2002, 2004). The large pedestal in the ‘Non-matched’ condition, which generally extended beyond the target area, might affect performance and enable learning in a way similar to that of the added flankers. The lack of long-term improvement across days in the ‘Matched’ condition is compatible with some previous studies (Dorais & Sagi, 1997), but not with all (Yu et al., 2004). Contrast discrimination learning is usually stronger with longer training periods and is diminished with roving stimuli (Adini et al., 2004; Yu et al., 2004). Here we argue that performance on the task is improved by adaptation adjusting gain control mechanisms or noise mechanisms that are dependent on contrast. Therefore, roving between multiple base contrasts and sizes might interfere with the contrast-dependent adjustments of previously adapted contrasts, resulting in diminished learning. This is in agreement with the Teich & Qian (2003) claim that adaptation to behaviorally relevant stimuli may be viewed as a short-term form of learning. According to their recurrent model, adaptation induces short-term changes that gradually become permanent with experience (Teich & Qian, 2003).

## Acknowledgments

This work was supported by the Basic Research Foundation administered by the Israel Academy of Sciences and Humanities (grant No. 6501560). We thank Dr. Misha Katkov for helpful comments.

## Appendix A Practicing CD with the target subtracted from the pedestal

## Stimuli and procedure

This pilot experiment was very similar to our main experiment. The main difference was that here the target was subtracted from, instead of added to, the pedestal. Thus, observers reported which of the Gabor patterns (left or right) appeared to have a lower contrast. However, there were some minor differences in the stimuli used. The pedestal Gabor had a phase of 90° and a constant base contrast of 63%, and the target Gabor had a phase of 270°. Eleven observers participated in this experiment. Five of them participated in the ‘Non-matched’ condition, and six participated in the ‘Matched’ condition. The pilot’ included five daily sessions, with three blocks in each session. The target contrast was determined by a staircase method, in which the contrast of the target decreased by 0.1 log units after three consecutive correct responses and increased by 0.1 log units after each mistake. This staircase converges at a level of 79.3% correct (Levitt, 1971). A block consisted of four parallel randomly interleaved staircases, one for each target size. For each staircase run, the threshold was calculated as the geometrical mean of all reversals except for the first three reversals, which were ignored. All staircases started with a high target contrast that enabled error-free detection or discrimination. In the ‘Non-matched’ condition the initial contrast was 31% for target sizes 0.37° and 0.56°, and 24% for target sizes 0.84° and 1.3°. In the ‘Matched’ condition the contrast was 24% for all target sizes.

## Results

Results for the average contrast discrimination thresholds (across observers) for the ‘Non-matched’ group and the ‘Matched’ group are displayed in Figures A1 and A2, respectively. To test for within-session improvements, we ran a repeated measures ANOVA with the within-observer factors including day (5 days), size (4 sizes), and block (first block vs last block), performed separately for the two groups. Performance within-session improved significantly in both groups, as indicated by a significant decrease in the threshold from the first block to the last block (11.8±1.9, F(1,4) = 39.8, p< 0.01; 1.3±0.5, F(1,5) = 7.7, p <0.05, mean±SEM in the ‘Non-matched’ group and the ‘Matched’ group, respectively). In the ‘Non-matched’ group, there was also a significant interaction between the block threshold and size (F(3,4) = 4.8, p< 0.05). To determine how the within-session improvements changed between sizes, we ran separate ANOVAs for each size. Results for the ‘Non-matched’ group show significant within-session improvements for every target size (16.3±1.7, F(1,4)=92.3, p=0.001, 15.6±4.4, F(1,4)=12.8, p<0.05, 8.0±2.2, F(1,4)=13.4, p<0.05, 7.1±1.5, F(1,4)=23.7, p<0.01, improvements in the 1^st^, 2^nd^, 3^rd^, and 4^th^ target sizes, respectively). However, in the ‘Matched’ group, only the two largest target sizes (0.84° and 1.3°) reached a significant within-session improvement (0.6±1.0, F(1,5)=0.4, p=0.6; 1.7±0.9, F(1,5)=3.4, p=0.1; 1.1±0.3, F(1,5)=13.7, p<0.05; 1.8±0.4, F(1,5)=18.6, p<0.01, improvements in the 1^st^, 2^nd^, 3^rd^, and 4^th^ target sizes, respectively).

Figure A suggests that many of the within-session gains, especially in the ‘Non-matched’ group, did not survive the time between sessions: the first measured threshold during a session was higher than the last measured threshold in a previous session. To further test the perseverance of the within-session improvements, we ran a repeated measures ANOVA with the within-observer factors of day (day 2-day 5), size (4 sizes), and block (the last block of the previous session vs the first block of the next session). Performance significantly deteriorated from the end of the previous session to the beginning of the next session only in the ‘Non-matched’ group (9.7±0.7, F(1,4)=213.6, p<0.001; 0.7±0.6, F(1,5)=1.2, p=0.3 for the ‘Non-matched’ group and the ‘Matched’ group, respectively), showing that the within-session gains were not fully retained between sessions. In the ‘Non-matched’ group, there was also a significant interaction between block threshold and size (F(3,4) = 3.9, p< 0.05). To determine how the between-session deterioration changed between sizes, we ran separate ANOVAs for each size. Results for the ‘Non-matched’ group show a significant between-session deterioration for every target size (13.4±2.6, F(1,4)=25.8, p=0.01, 14.1±3.0, F(1,4)=21.9, p<0.01, 6.3±1.7, F(1,4)=13.9, p<0.05, 5.0±0.4, F(1,4)=173.0, p<0.001 for deterioration in the 1^st^, 2^nd^, 3^rd^, and 4^th^ target sizes, respectively). In the ‘Matched’ group, none of the target sizes reached significant between-session deterioration (0.6±1.2, F(1,5)=0.28, p=0.6; 0.6±0.9, F(1,5)=0.4, p=0.5; 0.9±0.5, F(1,5)=4.2, p=0.1; 1.4±0.7, F(1,5)=3.9, p=0.1 for deterioration in the 1^st^, 2^nd^, 3^rd^, and 4^th^ target sizes, respectively). In addition, in both groups, performance was significantly enhanced when the size of the target increased (F(3,4) = 252.5, p<0.0001, F(3,5) = 3.4, p<0.05 in the ‘Non-matched’ group and the ‘Matched’ group, respectively).

Next, we tested for between-session improvements by running a repeated measures ANOVA with only the thresholds of the first day (day1) and the last day (day 5). The change in the overall daily performance (including all blocks of trials in a daily session) from the first session to the last session was significant only in the ‘Non-matched’ group (8.5±3.1, F(1,4) = 7.3, p=0.05, 2.2±1.3, F(1,5) = 2.9, p=0.2 in the ‘Non-matched’ group and the ‘Matched’ group, respectively).When testing for learning effects separately for the thresholds at the start (block 1) and at the end (block 3), there were no significant effects (start: 14.6±8.2, F(1,4) = 3.1, p =0.1, 3.2±1.8, F(1,5) = 3.3, p =0.1; end: 2.6±1.7, F(1,4) = 2.3, p =0.2, 1.3±0.9, F(1,5) = 2.5, p =0.2 in the ‘Non-matched’ group and the ‘Matched’ group, respectively).

To compare the performances of the groups, we added to the repeated measures ANOVAs the between-observer factor of group type (‘Non-matched’ group or the ‘Matched’ group). The improvements within-session (F(1,9) = 37.4, p< 0.001), the learning between days (first day vs last day, including all blocks in a session) (F(1,9) = 5.8, p< 0.05), and the deterioration between sessions (F(1,9) = 85.2, p< 0.0001) were significantly higher for the ‘Non-matched’ group. There was also a significant interaction between block threshold and size, indicating that the difference between groups in the within-session improvement depended on the size of the target (F(3,9) = 5.9, p< 0.01). Running separate ANOVAs for each size showed that for each size the within-session improvement was significantly higher for the ‘Non-matched’ group (F(1,9)=69.8, p<0.001; F(1,9)=27.1, p=0.001; F(1,9)=11.6, p<0.01; F(1,9)=17.6, p<0.01, improvements in the 1^st^, 2^nd^, 3^rd^, and 4^th^ target sizes, respectively). The difference between thresholds for different target sizes was also significantly lower for the ‘Matched’ group (F(3,9) = 195.5, p<0.0001). Table 1 summarizes the statistical results.

**Table A.** Statistical results for Exp1, ‘Matched’ vs ‘Non-matched’ conditions. *P < 0.05, **P < 0.01, ***P < 0.001.

| Factor | 'Non-matched' group | 'Matched' group | Between groups |
| --- | --- | --- | --- |
| Improvements within-session | ** | * | *** |
| Deteriorations between the end of the previous session and the beginning of the next session | *** | n.s | *** |
| Learning between the first day and the last day (all blocks) | * | n.s | * |
| Learning between the first day and the last day (the first block) | n.s | n.s | n.s |
| Learning between the first day and the last day (the last block) | n.s | n.s | n.s |
| Size | *** | * | *** |

**Fig. A.**
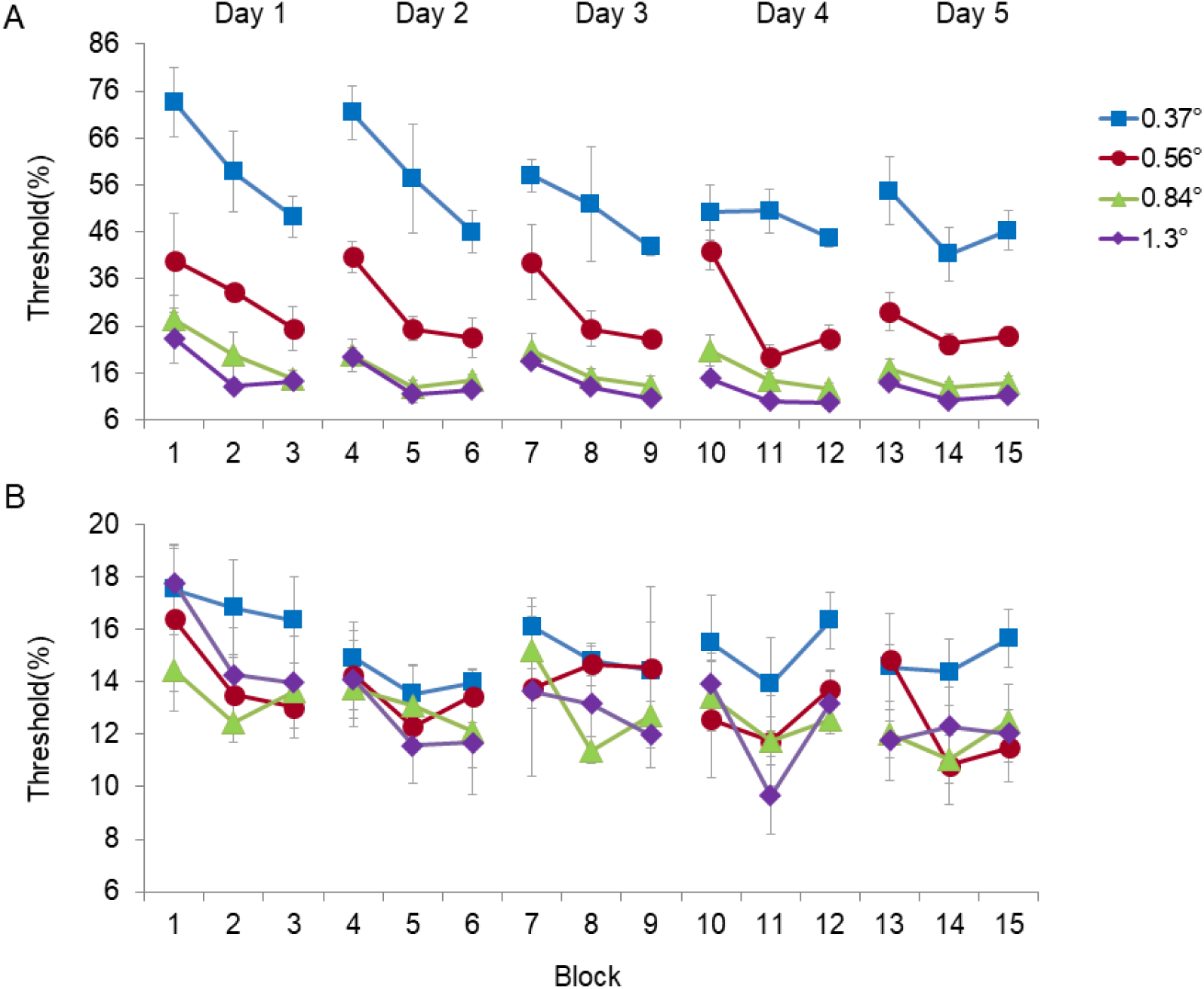
Contrast discrimination thresholds for stimulus configurations in which a target Gabor pattern (with sizes: 0.37°, 0.56°, 0.84°, or 1.3°) was subtracted from a pedestal Gabor pattern that was either fixed at a maximal size (1, ‘Non-matched’ group), or it had a size equal to the target size (2, ‘Matched’ group). Results show within-day improvements that are not fully retained in subsequent daily sessions. Thresholds are averages across observers (1: N = 5, 2: N=6), with error bars corresponding to ±1 standard error of the mean.

